# The Delta variant of SARS-CoV-2 maintains high sensitivity to interferons in human lung cells

**DOI:** 10.1101/2021.11.16.468777

**Authors:** Rayhane Nchioua, Annika Schundner, Susanne Klute, Sabrina Noettger, Fabian Zech, Lennart Koepke, Alexander Graf, Stefan Krebs, Helmut Blum, Dorota Kmiec, Manfred Frick, Frank Kirchhoff, Konstantin M.J. Sparrer

## Abstract

Interferons are a major part of the anti-viral innate defense system. Successful pathogens, including the severe acute respiratory syndrome coronavirus 2 (SARS-CoV-2), need to overcome these defenses to establish an infection. Early induction of interferons (IFNs) protects against severe coronavirus disease 2019 (COVID-19). In line with this, SARS-CoV-2 is inhibited by IFNs *in vitro*, and IFN-based therapies against COVID-19 are investigated in clinical trials. However, SARS-CoV-2 continues to adapt to the human population resulting in the emergence of variants characterized by increased transmission fitness and/or decreased sensitivity to preventive or therapeutic measures. It has been suggested that the efficient spread of these so-called “Variants of Concern” (VOCs) may also involve reduced sensitivity to IFNs. Here, we examined whether the four current VOCs (Alpha, Beta, Gamma and Delta) differ in replication efficiency or IFN sensitivity from an early isolate of SARS-CoV-2. All viruses replicated in a human lung cell line and in iPSC-derived alveolar type II cells (iAT2). The Delta variant showed accelerated replication kinetics and higher infectious virus production compared to the early 2020 isolate. Replication of all SARS-CoV-2 VOCs was reduced in the presence of exogenous type I, II and III IFNs. On average, the Alpha variant was the least susceptible to IFNs and the Alpha, Beta and Gamma variants show increased resistance against type III IFN. Although the Delta variant has outcompeted all other variants in humans it remained as sensitive to IFNs as an early 2020 SARS-CoV-2 isolate. This suggests that increased replication fitness rather than IFN resistance may be a reason for its dominance. Our results may help to understand changes in innate immune susceptibility of VOCs, and inform clinical trials exploring IFN-based COVID-19 therapies.

## Introduction

The IFN system constitutes a potent barrier against viral infections [1–3]. After recognition of viral pathogen-associated molecular patterns by germ-line encoded pattern recognition receptors, signaling cascades are activated. This results in the induction and secretion of IFNs as well as other pro-inflammatory cytokines [1]. The secreted IFNs can act in an autocrine or paracrine fashion. Upon binding to their respective receptors, the expression of hundreds of so-called interferon stimulated genes (ISGs) is induced, among them many well-known anti-viral factors [4,5]. IFNs are classified into three major types, based on the type of their receptor [6]. Human type I IFNs, including 13 subtypes of IFNα and IFNβ, bind to the IFNα/β receptor (IFNAR). Type II IFNs, such as IFNγ, interact with the IFN-gamma receptor (IFNGR). The Type III IFN family comprises four members (IFNλ1-4) which act via a complex composed of the Interleukin 10 receptor β-subunit (IL10R2) and the Interleukin 28 receptor α-subunit (IFNLR1). Type I and III IFNs may be secreted by almost any cell type, whereas type II IFN production is restricted to immune cells, particularly T and Natural Killer (NK) cells. Innate immunity plays a major role in defending against emerging pathogens like SARS-CoV-2, the causative agent of COVID-19 [7–11]. This respiratory virus has a profound global impact both on a socioeconomic level and as a major threat to human health. To date (November 4th, 2021), more than 240 million SARS-CoV-2 infections were reported worldwide, resulting in over 4.9 million deaths.

To be able to replicate in the presence of a functioning innate immune system, SARS-CoV-2 utilizes more than half of its about 30 proteins to suppress IFN induction and signaling [12–15]. However, despite these evasion mechanisms SARS-CoV-2 still remains sensitive towards all types of IFNs, with types II and III being most effective [12,16–19]. Importantly, early induction of high levels of IFNs in patients were reported to prevent severe COVID-19 [8,9]. Conversely, inborn defects in the IFN system or auto-antibodies against type I IFNs are frequently associated with severe disease [10,11]. Thus, IFN is currently evaluated in clinical trials as a therapeutic approach [20].

The efficient worldwide spread of the virus was associated with the emergence of novel, fitter variants that may show an increased ability to avoid innate immune control[21–23]. Currently, there are four recognized SARS-CoV-2 “Variants of Concern” (VOCs): B.1.1.7, B.1.351, P.1 and B.1.617.2 (World Health Organization, 2021). For simplification, these are also referred to as Alpha, Beta, Gamma and Delta variants, respectively. The order of their appearance in the human population is Beta, Alpha, Gamma and Delta. While the population spread of the Beta and Gamma variants was limited to certain regions, the Alpha variant rapidly overtook other strains in most countries in early 2021. However, within months it was outcompeted by the Delta VOC, which is currently (November 2021) responsible for about 90% of all SARS-CoV-2 infections worldwide. The fact that emerging VOCs outcompeted the early pandemic SARS-CoV-2 strains proves their increased fitness. A major hallmark of VOCs is increased escape from neutralizing antibodies [21,24]. However, VOCs also show various alterations outside the region encoding the Spike (S) protein that is the main target of the adaptive humoral immune response. Currently, it is poorly understood whether they also evolved increased resistance towards innate immune defenses.

## Results and Discussion

Next-generation sequencing of an early SARS-CoV-2 isolate from February 2020 (NL-02-2020) and four VOC isolates revealed amino acid changes in the S glycoprotein, as well as in proteins involved in replication and innate immune escape, compared to the first available sequence of the Wuhan-Hu-1isolate (Fig 1A). The impact of mutations in the S protein of VOCs has been the focus of many studies. Some of them are known to affect the affinity between S and the cellular receptor ACE2 and/or alter proteolytic activation of S, resulting in increased infectivity. In addition, mutations in E484, S477 or L452 in S allow the virus to evade adaptive immune responses [21,22,24]. Thus, for example the Delta variant was reported to be resistant to neutralization by some monoclonal antibodies and less susceptible towards patient sera [21]. However, the impact of alterations outside of S is poorly understood. The Delta VOC has accumulated 29 amino acid changes or deletions outside of the S region compared to the Wuhan-Hu-1 reference strain (Fig. 1A). These include mutations in Nsp3, ORF6 and N, which were reported to be crucial for IFN escape [12,25,26]. The consequence of these alterations is, however, unknown.

**Fig 1.**
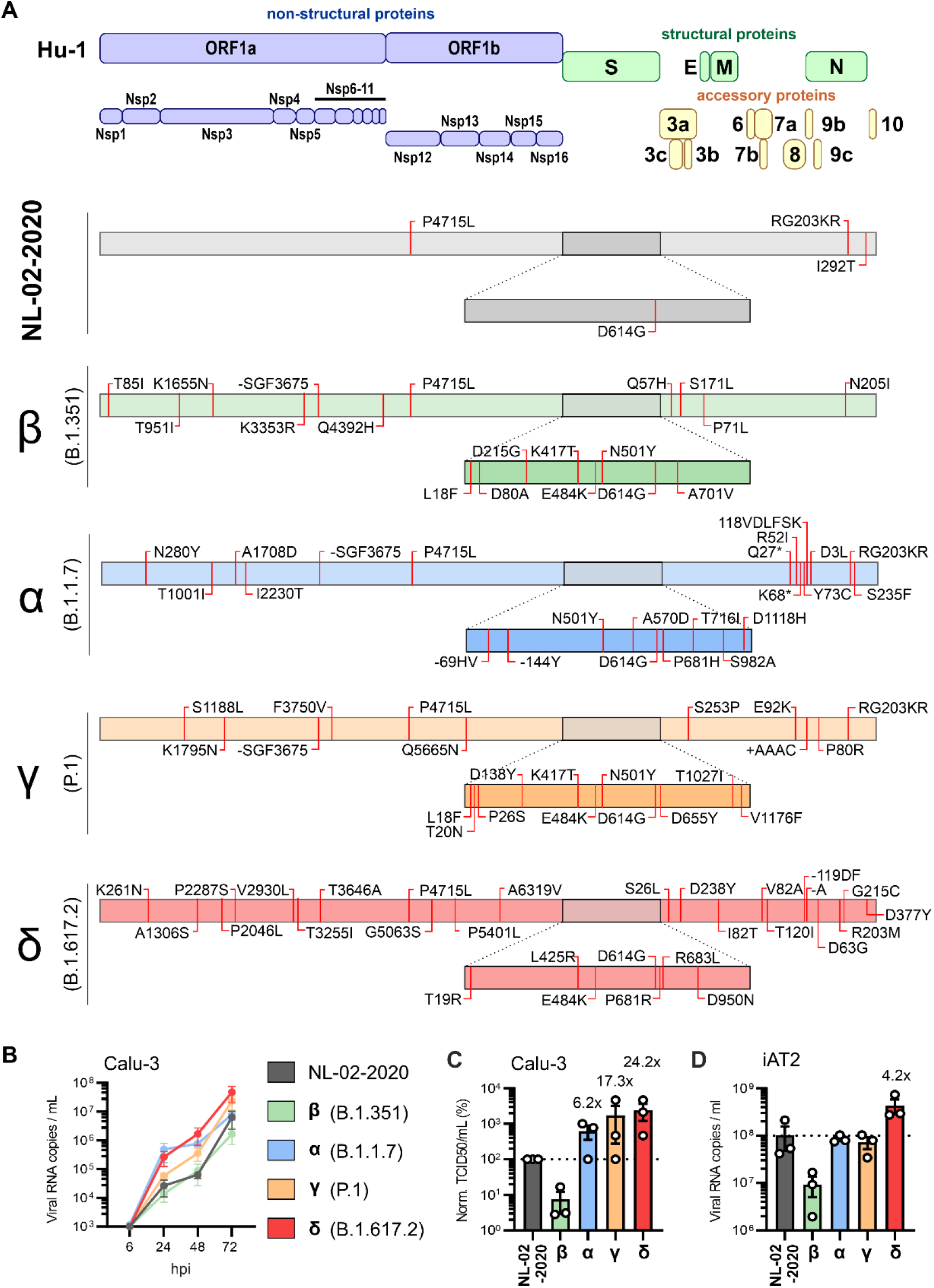
Amino acid differences and replication kinetics of early SARS-CoV-2 and VOCs. **A**, Schematic depiction of the SARS-CoV-2 genomic arrangement and proteins (top). Outline of the specific amino acid exchanges compared to the reference Hu-1 sequence in an early European Feb 2020 SARS-CoV-2 isolate (NL-02-2020), and four variants of concern in the order of appearance: Beta (B.1.351), Alpha (B.1.1.7), Gamma (P.1) and Delta (B.1.617.2) as assessed by next-generation sequencing assembly of the full genome. **B**, Viral RNA in the supernatant of Calu-3 cells infected with indicated SARS-CoV-2 variants was quantified by qRT-PCR at indicated timepoints post infection (MOI 0.05). Day 0 wash CTRL values were subtracted from data shown in the panel. n=3±SEM. **C**, Infectious SARS-CoV-2 particles in the supernatant, corresponding to the 72h post-infection time point shown in (B). n=3±SEM. **D**, Viral RNA in the supernatant of iAT2 cells infected with indicated SARS-CoV-2 variants was quantified by qRT-PCR at 2 days post infection (MOI 0.5) day 0 wash control values were subtracted from data shown in the panel. n=3±SEM. Related to Fig S1.

Cells of the respiratory tract are primary targets of SARS-CoV-2 infection. We found that the Delta variant replicated with higher efficiency (Fig 1B, Figs S1A and S1B) and produced ∼24-fold higher infectious titers at 3 days post-infection (Fig 1C) compared to the NL-02-2020 isolate in the human lung cell line Calu-3. The Alpha and Gamma variants showed intermediate phenotypes, while the Beta variant replicated with moderately reduced efficiency compared to NL-02-2020. The Delta VOC replicated with ∼4-fold higher efficiency than all other SARS-CoV-2 isolates analyzed in iPSC-derived alveolar epithelial type II (iAT2) cells (Fig 1D). AT2 cells constitute approximately 60% of the pulmonary alveolar epithelial cells and are the main targets of SARS-CoV2 in the distal lung [27]. Virus-induced loss of AT2 cells is linked to the severity of COVID-19 associated acute respiratory distress syndrome [28] and reduced lung regeneration [27]. Altogether, the Delta variant that currently dominates the COVID-19 pandemic clearly showed the highest replication efficiencies in human lung cells.

Consistent with published results [12,29], the NL-02-2020 isolate was more sensitive towards IFNβ, IFNγ and IFNλ1 than to IFNα2 in Calu-3 cells (Figs 2A and 2B and Fig S2A). IFN treatment did not affect cell viability (Fig S2B). All VOCs were still susceptible towards exogenous IFNs, albeit not to the same extent as NL-02-20 (Figs 2A and 2B). Area under the curve analysis (Fig 2B) as an indicator of virus production in the presence of different concentrations of IFNs revealed that the Beta and Gamma VOC showed moderately increased and the Alpha variant the highest (8-fold) resistance against IFN treatment. Susceptibility of the Alpha, Delta and especially the Gamma variant towards IFNα2 was 3 to 10-fold decreased. The Beta, Alpha and Gamma variants were approximately 5-fold less susceptible towards IFNλ1 than the NL-02-2020 and Delta isolates (Fig 2C). Notably, the Delta variant remained overall at least as sensitive towards IFN treatment as the NL-02-2020 isolate (Figs 2B and 2C).

**Fig 2.**
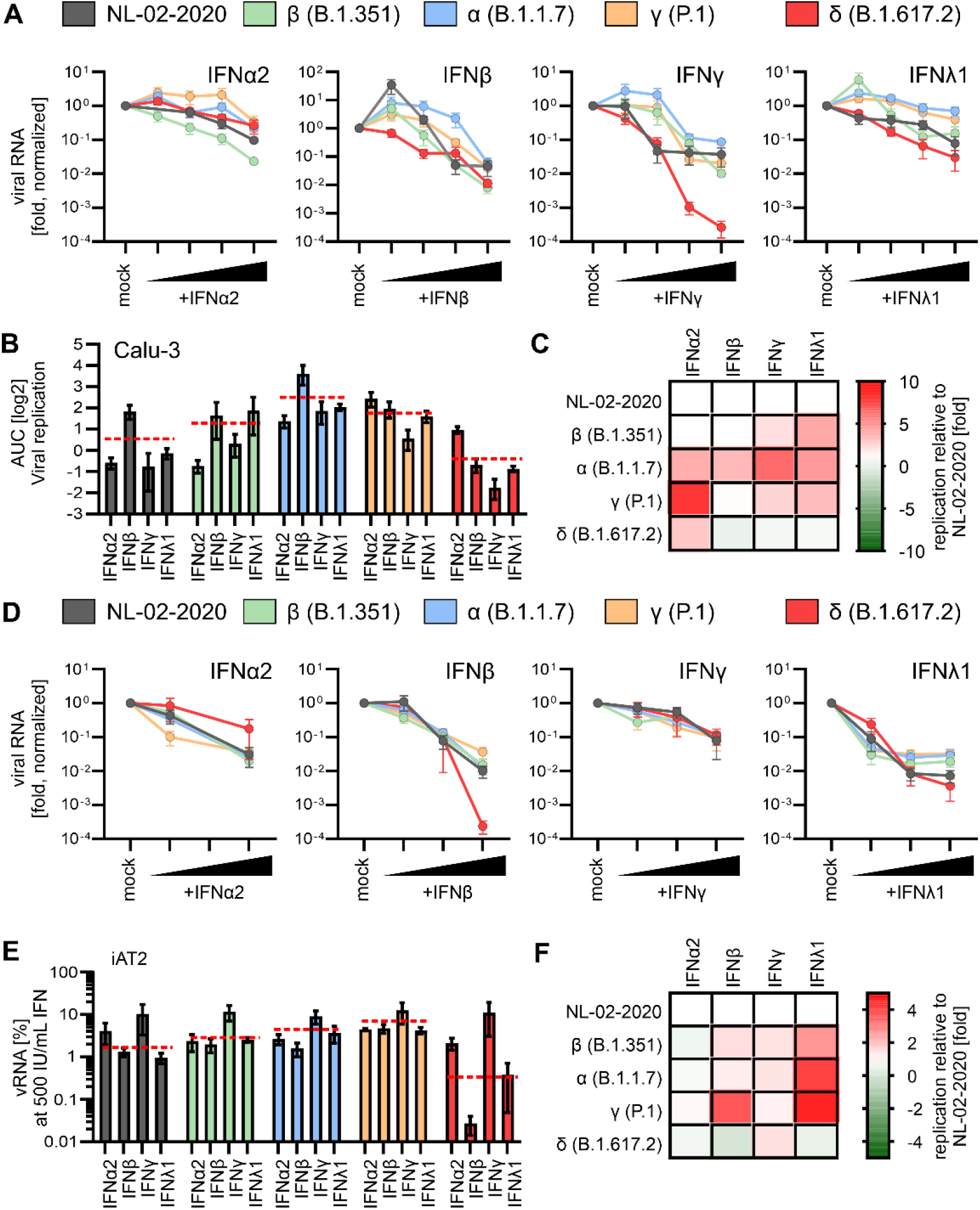
Interferon sensitivity of NL-02-2020 and VOCs. **A**, Normalized amount of viral RNA in the supernatant of Calu-3 cells infected with indicated SARS-CoV-2 variants was quantified by qRT-PCR at 72h post-infection (MOI 0.05, no IFN set to 100%). Cells were infected 3 days post treatment with indicated IFNs (α2, β and γ 0.5, 5, 50 and 500 U/ml) or IFNλ1 (0.1, 1, 10 and 100 ng/ml). n=2±SEM. **B**, Area under the curve analysis of the data in (A) representing the replication of the variants in the presence of IFNs. Red lines indicate the average over all IFNs for one variant. **C**, Heatmap displaying differences in viral replication (Area under the curve analysis) of the VOCs compared to the NL-02-2020 variant upon IFN treatment of Calu-3 cells. Red, increased replication, green, decreased replication relative to replication of NL-02-2020. Data from (A). **D**, Normalized amount of viral RNA in the supernatant of iAT2 cells infected with indicated SARS-CoV-2 variants as quantified by qRT-PCR at 48h post-infection (MOI 0.5, no IFN set to 100%). Cells were infected for 2 days post treatment with indicated IFNs (α2, β and γ: 5, 50 and 500 U/ml) or IFNλ1 (1, 10 and 100 ng/ml). n=4±SEM. **E**, Percentage of viral RNA in the supernatant of iAT2 cells as a fraction between non-treated and IFN treated (500 IU/mL or 100 ng/mL). data from (D). Red lines indicate the average over all IFNs for one variant. **F**, Heatmap displaying fold differences in viral replication of the VOCs in iAT2 cells compared to the NL-02-2020 variant (set to 1) upon treatment with IFN (500 IU/mL or 100 ng/mL). Red, increased replication, green, decreased replication relative to NL-02-2020. Data from (D). Related to Figs S2 and S3.

For the most part, analysis of iAT2 cells confirmed the results obtained in Calu-3 cells (Figs 2D-F, S3A). However, type II IFN was less effective against SARS-CoV-2 in iAT2 cells than in Calu-3 cells, suggesting different receptor expression or pathway activity (Fig. 2D). In comparison, IFNβ and IFNλ1 remained highly efficient and decreased viral RNA levels by more than two orders of magnitude. Controls showed the metabolic activity of iAT2 cells was not affected by IFN treatment (Fig S3B). Focusing on the viral load at the highest IFN dose compared to the non-treated sample revealed all VOCs remained sensitive towards IFN treatment (Figs 2E and 2F), with the Gamma and Alpha variant being ∼3- and 4-fold less affected by IFN treatment, respectively. In comparison to the early NL-02-2020 isolate the Beta, Alpha and Gamma VOCs were 3-5-fold less susceptible to inhibition by type III IFN (Fig 2F). In line with the results in Calu-3 cells, the Delta variant was still sensitive towards IFN, even more so in the case of IFNβ.

A variety of viral features, including virion infectivity, replication efficiency, and efficiency of immune evasion, may contribute to the emergence of SARS-CoV-2 VOCs. Antibody escape of VOCs has been extensively studied [30–32] and especially the Delta variant showed decreased sensitivity to neutralization [21]. Here, we confirmed [23] that the Alpha variant shows reduced susceptibility to IFNs. However, this VOC showed little if any increase in replication fitness in human lung cells. In comparison, the dominating Delta variant clearly replicates to higher levels but remains highly susceptible to inhibition by IFNs. It is tempting to speculate that the Alpha variant had an advantage over the original SARS-CoV-2 strains because of its reduced IFN sensitivity but was later outcompeted due to the increased replication fitness/transmission of the Delta variant. Coronaviruses are prone to recombination and combination of the IFN resistance of the Alpha variant with the replication advantage of the Delta variant poses a threat for the emergence of a new VOC.

Reduced susceptibility towards IFNs is likely determined by mutations outside of S. Notably, the Alpha and Gamma VOCs showing reduced type III IFN sensitivity share a deletion in Nsp6, which was previously implicated in inhibiting innate immune responses [12,14]. SARS-CoV-2 utilizes most of its genes to suppress or counteract innate immune defense mechanisms [12,14]. Thus, these viral countermeasures most likely play a key role in virus transmission. Further studies on the molecular determinants of reduced IFN sensitivity and improved innate immune evasion of emerging SARS-CoV-2 variants are highly warranted.

Our results indicate that IFNβ and IFNλ1 are most effective in inhibiting the Delta VOC in human alveolar epithelial type II cells proposed to play a key role in the spread and pathogenesis of SARS-CoV-2. This finding might help to improve IFN-based therapies against the SARS-CoV-2 VOC that currently dominates the pandemic.

## Materials and Methods

### Cell culture

Calu-3 (human epithelial lung adenocarcinoma) cells were cultured in Minimum Essential Medium Eagle (MEM, Sigma, Cat#M4655) supplemented with 10% (upon and after viral infection) or 20% (during all other times) heat-inactivated fetal bovine serum (FBS, Gibco, Cat#10270106), 100 units/ml penicillin, 100 µg/ml streptomycin (ThermoFisher, Cat#15140122), 1 mM sodium pyruvate (Pan Biotech, Cat#P04-8010), and 1x non-essential amino acids (Sigma, Cat#M7145). Vero E6 cells (*Cercopithecus aethiops* derived epithelial kidney, ATCC) and TMPRSS2-expressing Vero E6 cells (kindly provided by the National Institute for Biological Standards and Control (NIBSC), No. 100978) were grown in Dulbecco’s modified Eagle’s medium (DMEM, Gibco, Cat#41965039) supplemented with 2.5% (upon and after viral infection) or 10% (during all other times) heat-inactivated FBS (Gibco, Cat#10270106), 100 units/ml penicillin, 100 µg/ml streptomycin (ThermoFisher, Cat#15140122), 2 mM L-glutamine (Gibco, Cat#25030081), 1 mM sodium pyruvate (Pan Biotech, Cat# P04-8010), 1x non-essential amino acids (Sigma, Cat#M7145) and 1 mg/mL Geneticin (Gibco, Cat#10131-019) (for TMPRSS2-expressing Vero E6 cells). Caco-2 cells (human epithelial colorectal adenocarcinoma, kindly provided by Prof. Holger Barth (Ulm University)) were grown in the same media as Vero E6 cells but with supplementation of 10% heat-inactivated FBS.

Human induced Alveolar Type 2 cells (iAT2) were differentiated from BU3 NKX2-1^GFP^;SFTPC^tdTomato^ induced pluripotent stem cells [33] (iPCSs, kindly provided by Darrell Kotton, Boston University and Boston Medical Center) and maintained as alveolospheres embedded in 3D Matrigel in CK+DCI media, as previously described [34]. For infection studies, iAT2 cells were cultured as 2D cultures on Matrigel-coated plates in CK+DCI medium + 10 µM Y-27632 (Tocris, Cat#1254) for 48 h before switching to CK+DCI medium on day 3.

### SARS-CoV-2 stocks

The SARS-CoV-2 variant B.1.351 (Beta), 2102-cov-IM-r1-164 [35] was provided by Prof. Michael Schindler (University of Tübingen) and the B.1.617.2 (Delta) variant by Prof. Florian Schmidt (University of Bonn). The BetaCoV/Netherlands/01/NL/2020 (NL-02-2020) and B.1.1.7. (Alpha) variants were obtained from the European Virus Archive. The hCoV-19/Japan/TY7-503/2021 (Brazil P.1) (Gamma) (#NR-54982) isolate was obtained from the BEI resources. SARS-CoV-2 strains were propagated on Vero E6 (NL-02-2020, Delta), VeroE6 overexpressing TMPRSS2 (Alpha), CaCo-2 (Beta) or Calu-3 (Gamma) cells. To this end, 70-90% confluent cells in 75 cm^2^ cell culture flasks were inoculated with the SARS-CoV-2 isolate (multiplicity of infection (MOI) of 0.03-0.1) in 3.5 ml serum-free medium. The cells were incubated for 2h at 37°C, before adding 20 ml medium containing 15 mM HEPES (Carl Roth, Cat#6763.1). Virus stocks were harvested as soon as strong cytopathic effect (CPE) became apparent. The virus stocks were centrifuged for 5 min at 1,000 g to remove cellular debris, aliquoted, and stored at -80°C until further use.

### Sequencing of full-length SARS-CoV-2 genomes

Virus stocks were inactivated and lysed by adding 0.3 ml TRIzol Reagent (ambion, Cat#132903) to 0.1 ml virus stock. Viral RNA was isolated using the Direct-zol RNA MiniPrep kit (ZymoResearch, Cat#R2050) according to manufacturer’s instructions, eluting the RNA in 50 µl DNase/RNase free water. The protocol to prepare the viral RNA for sequencing was modified from the nCoV-2019 sequencing protocol V.1. For reverse transcription, the SuperScript IV First-Strand Synthesis System (Invitrogen, Cat#18091050) was used with modified manufacturer’s instructions. First, 1 µl random hexamers (50 ng/µl), 1 µl dNTPs mix (10 mM each), and 11 µl template RNA (diluted 1:10 in DNase/RNase free water) were mixed, incubated at 65°C for 5 min and placed on ice for 1 min. Next, 4 µl SSIV Buffer, 1 µl DTT (100 mM), 1 µl RNaseOUT RNase Inhibitor, and 1 µl SSIV Reverse Transcriptase were added to the mix, followed by incubation at 24°C for 5 min, 42°C for 50 min, and 70°C for 10 min. To generate 400 nt fragments in PCR, the ARTIC nCoV-2019 V3 Primer set (IDT) and the Q5 Hot Start High-Fidelity 2X Master Mix (NEB, Cat#M0494S) were used with modified manufacturer’s instructions. The primers pools 1 and 2 were diluted to a final concentration of 10 µM and a reaction with each primer pool was set up as follows, 4 µl respective primer pool, 12.5 µl Q5 Hot Start High-Fidelity 2X Master Mix, 6 µl water, and 2.5 µl cDNA. The PCR was performed as follows, 98°C for 30 s, 30 cycles of 98°C for 15 s and 65°C for 5 min, and hold at 4°C. The PCR products were run on a 1% agarose gel to check for the presence of fragments at the correct size. The products from primer pool 1 and primer pool 2 for each variant were pooled, diluted and quantified by Qubit DNA HS kit (Thermo Fisher, Cat# Q32851). The sequencing amplicon pools were diluted to 0.2 ng/µl and tagmented with Nextera XT library prep kit (Illumina, Cat#FC-131-1024). Nextera libraries were dual-barcoded and sequenced on an Illumina NextSeq1000 instrument. The obtained sequenced reads were demultiplexed and mapped against the SARS-CoV-2 reference genome (NC_045512.2) with BWA-MEM[36]. Pileup files were generated from the mapped reads using Samtools[37]. The mapped reads and the pileup file were used to construct the consensus sequence with the iVar package[38] using default settings.

### Plaque-forming Unit (PFU) assay

To determine the infectious titres, SARS-CoV-2 stocks were serially diluted 10-fold. Monolayers of Vero E6 cells in 12-wells were infected with the dilutions and incubated for 1 to 3 h at 37°C with shaking every 15 to 30 min. Afterwards, the cells were overlayed with 1.5 ml of 0.8 % Avicel RC-581 (FMC Corporation) in medium and incubated for 3 days. The cells were fixed by adding 1 ml 8 % paraformaldehyde (PFA, Sigma-Aldrich, Cat#158127-100G) in Dulbecco’s phosphate buffered saline (DPBS, Gibco, Cat#14190144) and incubated at room temperature for 45 min. After discarding the supernatant, the cells were washed with DPBS (Gibco, Cat#14190144) once, and 0.5 ml of staining solution (0.5% crystal violet (Carl Roth, Cat#42555) and 0.1% triton X-100 (Sigma-Aldrich, Cat#X100-100ML) in water) was added. After 20 min incubation at room temperature, the staining solution was removed using water, virus-induced plaque formation quantified, and plaque forming units per ml (PFU/ml) calculated.

### SARS-CoV-2 variants replication kinetics

1.5×10^5^ Calu-3 cells were seeded in 24-well plates. 24 h post-seeding, Calu-3 cells were infected with the different variants of SARS-CoV-2 (MOI 0.05). 5h later, supernatant was removed and 1 ml of fresh medium was added. 6h, 24h, 48h and 72h post-infection, supernatants were harvested for qRT-PCR analysis.

### Effect of IFNs on SARS-CoV-2 replication

1.5×10^5^ Calu-3 cells were seeded in 24-well plates. 24 h and 96 h post-seeding, cells were stimulated with increasing amounts of IFNs (α2 (R&D Systems, Cat#11101-2), β (R&D Systems, Cat#8499-IF-010/CF) and γ (R&D Systems, Cat#285-IF-100/CF). 0.5, 5, 50 and 500 U/ml) or IFN-λ1 (R&D Systems, Cat#1598-IL-025/CF) 0.1, 1, 10 and 100 ng/ml) in 0.5 ml of medium. 6 to 12 h after the first stimulation, the medium was replaced. 2 h after the second stimulation, Calu-3 cells were infected with the indicated SARS-CoV-2 variants (MOI 0.05) and 5 to 6h later, supernatant was removed, cells were washed once with DPBS (Gibco, Cat#14190144) and 0.5 ml fresh medium was added. 6 (wash control), 24, 48 and 72 h post-infection, supernatants were harvested for qRT-PCR analysis.

1.5×10^4^ – 3×10^4^ iAT2 cells were seeded as single cells in 96-well plates coated for 1 h at 37 °C with 0.16 mg/ml Matrigel (Corning, Cat#356238) diluted in DMEM/F12 (Thermo Fisher, Cat#11330032). 48h post-seeding, cells were stimulated with increasing amounts of IFNs (α2 (R&D Systems, Cat#11101-2), β (R&D Systems, Cat#8499-IF-010/CF) and γ (R&D Systems, Cat#285-IF-100/CF). 0.5, 5, 50 and 500 U/ml) or IFNλ1 (R&D Systems, Cat#1598-IL-025/CF) 0.1, 1, 10 and 100 ng/ml) in 150 µl medium. 24 h post-treatment, iAT2 cells were infected with the indicated SARS-CoV-2 variants (MOI 0.5). 5 to 6 h later, supernatants were removed, cells were washed once with DPBS (Gibco, Cat#14190144) and 200 µl of fresh medium was added. Supernatants were harvested at 6 h (wash control) and 48 h post-infection for qRT-PCR analysis.

### qRT-PCR

N (nucleoprotein) transcript levels were determined in supernatants collected from SARS-CoV-2 infected Calu-3 or iAT2 cells 6, 24, 48, and 72 h post-infection as previously described [39]. Total RNA was isolated using the Viral RNA Mini Kit (Qiagen, Cat#52906) according to the manufacturer’s instructions. qRT-PCR was performed as previously described [39] using TaqMan Fast Virus 1-Step Master Mix (Thermo Fisher, Cat#4444436) and a OneStepPlus Real-Time PCR System (96-well format, fast mode). Primers were purchased from Biomers (Ulm, Germany) and dissolved in RNase free water. Synthetic SARS-CoV-2-RNA (Twist Bioscience, Cat#102024) or RNA isolated from BetaCoV/France/IDF0372/2020 viral stocks quantified via this synthetic RNA (for low Ct samples) was used as a quantitative standard to obtain viral copy numbers. All reactions were run in duplicates. Forward primer (HKU-NF): 5’-TAA TCA GAC AAG GAA CTG ATT A-3’; Reverse primer (HKU-NR): 5’-CGA AGG TGT GAC TTC CAT G-3’; Probe (HKU-NP): 5’-FAM-GCA AAT TGT GCA ATT TGC GG-TAMRA-3’.

### Tissue Culture Infection Dose50 (TCID50) endpoint titration

SARS-CoV-2 stocks or infectious supernatants were serially diluted. 25,000 Caco-2 cells were seeded per well in 96 F-bottom plates in 100 µl medium and incubated overnight. Next, 50 µl of diluted SARS-CoV-2 stocks or infectious supernatants were used for infection, resulting in final dilutions of 1:101 to 1:1012 on the cells in nine technical replicates. Cells were then incubated for 5 days and monitored for CPE. TCID50/ml was calculated according to the Reed and Muench method.

### MTT (3-[4,5-dimethyl-2-thiazolyl]-2,5-diphemyl-2H-tetrazolium bromide) assay

6×10^4^ Calu-3 cells were seeded in 96-well F-bottom plates. 1.5×10^4^ – 3×10^4^ iAT2 cells were seeded as single cells in 96-well F-bottom plates, coated for 1h at 37°C with 0.16 mg/ml Matrigel (Corning, Cat#356238) diluted in DMEM/F12 (Thermo Fisher, Cat#11330032). The cells were stimulated with increasing amounts of IFNs (α2 (R&D Systems, Cat#11101-2), β (R&D Systems, Cat#8499-IF-010/CF) and γ (R&D Systems, Cat#285-IF-100/CF). 0.5, 5, 50 and 500 U/ml) or IFN-λ1 (R&D Systems, Cat#1598-IL-025/CF) 0.1, 1, 10 and 100 ng/ml) in 100 µl of medium 24 h and 96 h post-seeding. 6 h after the first stimulation, the medium was replaced. To analyze the cell viability of Calu-3 cells and iAT2 cells after interferon treatment, 100 µl of MTT solution (0.5 mg/ml in DPBS (Gibco, Cat#14190144)) was added to the cells 2 h after the second stimulation and the cells were incubated for 3 h at 37 °C. Subsequently, the supernatant was discarded and 100 µl 4% PFA (Sigma-Aldrich, Cat#158127-100G) in DPBS (Gibco, Cat#14190144) was added for 20 min. After washing with 100 µl DPBS (Gibco, Cat#14190144), the formazan crystals were dissolved in 100 µl of a 1:2 mixture of dimethyl sulfoxide (DMSO, Invitrogen, Cat#D12345) and ethanol. Absorption was measured at 490 nm with baseline corrected at 650 nm by using a Vmax kinetic microplate reader (Molecular Devices) and the SoftMax Pro 7.0.3 software.

## Acknowledgements

We thank Daniela Krnavek, Martha Mayer, Kerstin Regensburger, Regina Burger, Jana-Romana Fischer and Birgit Ott for laboratory assistance. We also thank Michael Schindler for providing the B.1.351 (Beta) variant and Florian Schmidt, Beate M. Kümmerer and Hendrick Streeck for providing the B.1.617.2 (Delta) variant. The following reagent was obtained through BEI Resources, NIAID, NIH: SARS-Related Coronavirus 2, Isolate hCoV-19/Japan/TY7-503/2021 (Brazil P.1), NR-54982, contributed by National Institute of Infectious Diseases. hCoV-19/Netherlands/01/NL/2020 and nCoV19 isolate/England/MIG457/2020 were provided via the European Virus archive. The BU3-NGST iPSC line was generated with funding from the National Center for Advancing Translational Sciences (“NCATS”) (grant U01TR001810) and kindly provided by D.N. Kotton, (Center for Regenerative Medicine, Boston Medical Center). This study was supported by DFG grants to F.K., K.M.J.S., D.K. and M.F. (F.K. and K.M.J.S., CRC 1279 and SPP1923; D.K., KM 5/1-1; M.F., 278012962 and 458685747), the BMBF to F.K. and K.M.J.S. (Restrict SARS-CoV-2 and IMMUNOMOD) and a COVID-19 research grant from the Ministry of Science, Research and the Arts of Baden-Württemberg (MWK) to F.K.. K.M.J.S. was additionally supported by a COVID-19 emergency grant from the DFG (SP1600/6-1). A joint project (Bay-VOC) from the Bavarian State Ministry of the Environment and Public Health supported A.G, S.K and H.B. L.K., S.K. and A.S. are part of the International Graduate School for Molecular Medicine, Ulm (IGradU).

## Author Contribution

Conceptualization and funding acquisition, F.K., K.M.J.S.; Investigation, R.N., S.K., A.S., F.Z.; iAT2 cells, A.S., M.F.; NGS preparation, sequencing and analysis, L.K., A.G., S.K. and H.B.; Writing, F.K. and K.M.J.S; Review and editing, all authors.

## Data Availability

The source data is available from the corresponding authors upon reasonable request. Full length sequences were deposited to GSAID (IDs will be added once approved).

## Supporting information

**S1 Fig.**
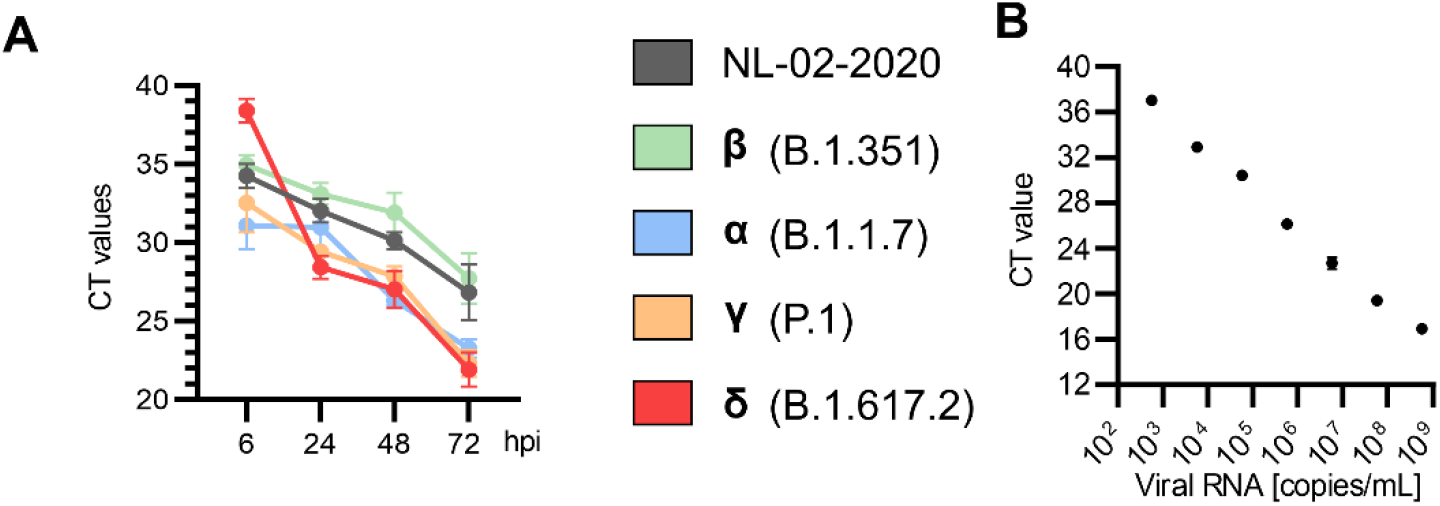
Replication kinetics of NL-02-2020 and VOCs in lung cells. **A**, Raw qRT-PCR Ct values obtained from supernatants of Calu-3 cells infected with indicated SARS-CoV-2 variants (MOI 0.05), at indicated time points. n=3±SEM. **B**, Exemplary standard curve used for Viral RNA loads quantification.

**S2 Fig.**
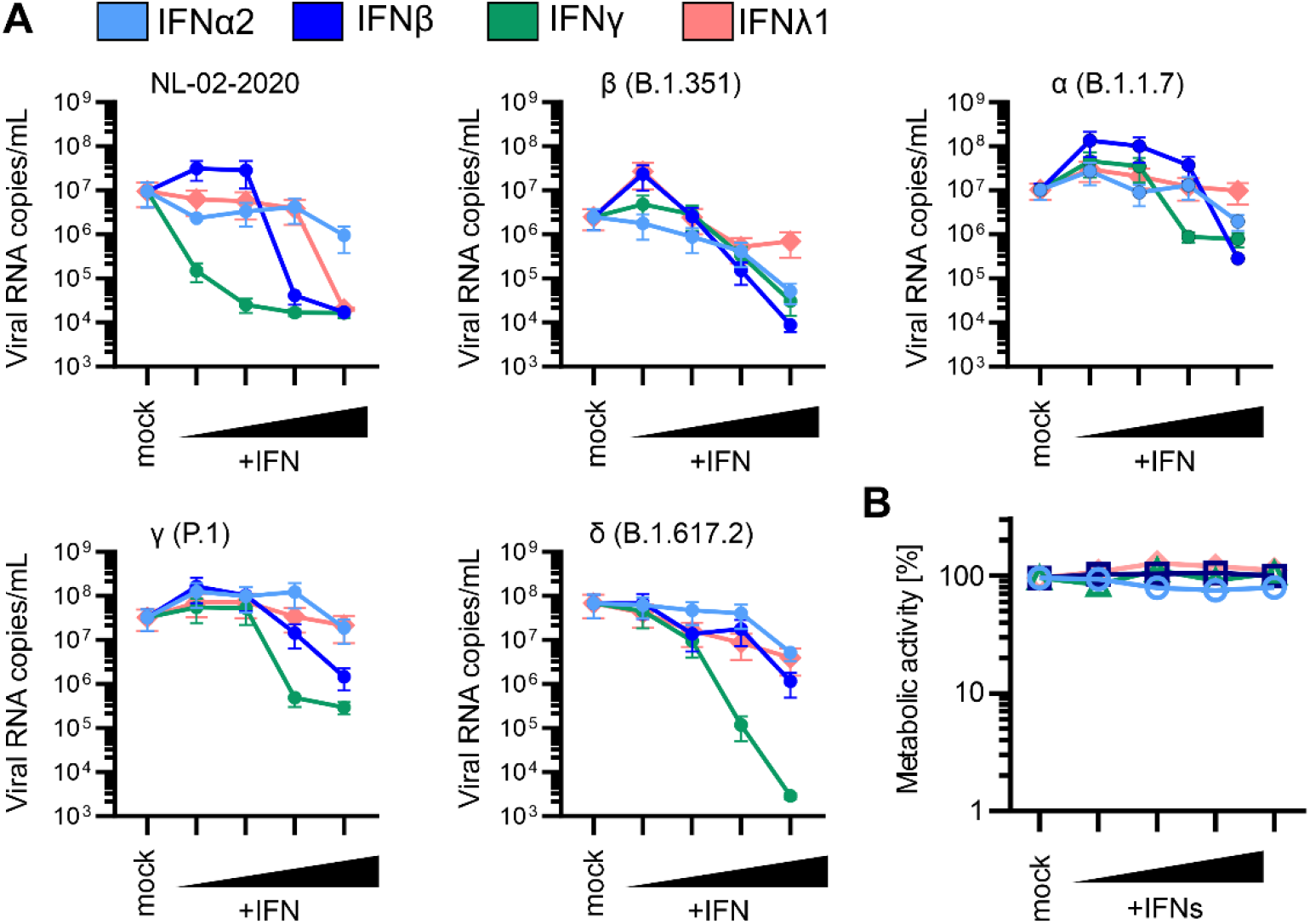
Interferon sensitivity of NL-02-2020 and VOCs in Calu-3 cells. **A**, Viral RNA levels in the supernatant of Calu-3 cells infected with indicated SARS-CoV-2 variants and quantified by qRT-PCR at 72h post-infection (MOI 0.05). n=2±SEM. **B**, Metabolic activity of Calu-3 cells after treatment with IFNs as in (A). n=3±SEM.

**S3 Fig.**
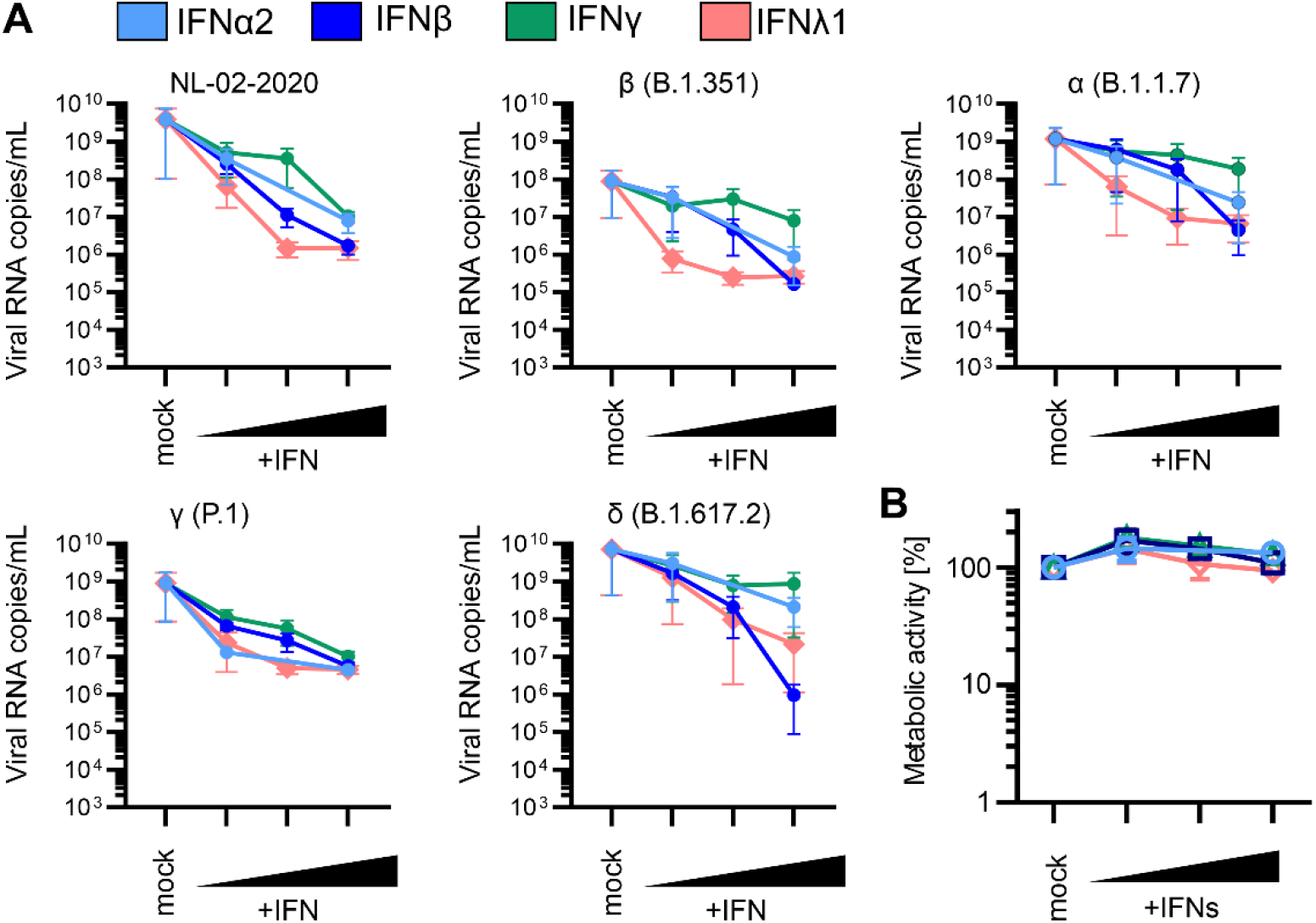
Interferon sensitivity of NL-02-2020 and VOCs in iAT2 cells. **A**, Viral RNA levels in the supernatant of iAT2 cells infected with indicated SARS-CoV-2 variants and quantified by qRT-PCR at 48h post-infection (MOI 0.5). (n=4±SEM). **B**, Metabolic activity of iAT2 cells after treatment with IFNs as indicated in panel (A). n=3±SEM.

